# Calcium Imaging Reveals Host-Graft Synaptic Network Formation in Spinal Cord Injury

**DOI:** 10.1101/795583

**Authors:** S Ceto, KJ Sekiguchi, Y Takashima, A Nimmerjahn, MH Tuszynski

## Abstract

Neural stem/progenitor cell grafts integrate into sites of spinal cord injury (SCI) and form anatomical and electrophysiological neuronal relays across lesions. To determine how grafts become synaptically organized and connect with host systems, we performed calcium imaging of neural progenitor cell grafts within sites of SCI, using both *in vivo* imaging and spinal cord slices. Stem cell grafts organize into localized synaptic networks that are spontaneously active. Following optogenetic stimulation of host corticospinal tract axons regenerating into grafts, distinct and segregated neuronal networks respond throughout the graft. Moreover, optogenetic stimulation of graft axons extending out from the lesion into the denervated spinal cord also trigger responses in local *host* neuronal networks. In vivo imaging reveals that behavioral stimulation of host elicits focal synaptic responses within grafts. Thus, remarkably, neural progenitor cell grafts form functional synaptic subnetworks in patterns paralleling the normal spinal cord.

## INTRODUCTION

Eliciting regeneration of the injured adult central nervous system (CNS) is, despite recent progress, extraordinarily difficult. While experimental approaches targeting multiple mechanisms can succeed in promoting injured host axonal regeneration into and even beyond sites of spinal cord injury (SCI) (Alto et al., 2009; Assinck et al., 2017; Bráz et al., 2012; Jayaprakash et al., 2019; Liu et al., 2010; Lu et al., 2012), the quantity and distance of host axonal regeneration is limited: typically, only hundreds of host axons regenerate for distances that rarely exceed 1-2 mm. In contrast, recent reports indicate that neural progenitor cells (NPCs) or neural stem cells (NSCs) implanted into sites of SCI support extensive host axon regeneration into grafts occupying the lesion site, while axons emerging from the grafted neurons extend into the distal host spinal cord in very large numbers (tens to hundreds of thousands of axons) and for long distances (up to 50 mm) (Kadoya et al., 2016; Lu et al., 2012, 2014; Rosenzweig et al., 2018). Indeed, after even complete spinal cord transection, NPC and NSC grafts support significant functional improvement (Koffler et al., 2019; Lu et al., 2012), and this functional improvement is abolished if the spinal cord is re-transected immediately rostral to the lesion site, suggesting that novel synaptic relays have formed from host-to-graft-to-host across the lesion site.

Little is known, however, regarding the synaptic architecture generated in NPC and NSC grafts placed in host spinal cord lesion sites. It is clear that host axons regenerating into grafts form synaptic contacts onto graft neurons (Adler et al., 2017; Dulin et al., 2018; Kumamaru et al., 2019; Lu et al., 2012; Rosenzweig et al., 2018) and are capable of generating post-synaptic currents (Kadoya et al., 2016). Similarly, graft neurons in sites of stroke in the cerebral cortex respond to optogenetic stimulation of thalamo-cortical axons as well as physiological tactile stimulation (Tornero et al., 2017). Graft neurons can also modulate activity in host neuron populations (Etlin et al., 2016; Steinbeck et al., 2015). However, how host axons regenerating into grafts influence graft activity has not been described, and several possibilities exist. First, regenerating host axons may form sparse functional connections with graft neurons that represent single isolated neuronal relays. Second, regenerating host axons may recruit larger networks of graft neurons that are similar to previously described targets of corticospinal projections in the intact spinal cord (Lemon, 2008; Levine et al., 2014; Ueno et al., 2018). A third possibility is that areas of dense axon regeneration act as initiation points of high frequency spiking activity that then activate neighboring populations of neurons; this would result in a slow spread of excitation across the graft, reminiscent of the slow waves of “depression” that occur in epilepsy, stroke and other neural disorders (Bikson et al., 2003; Pinto et al., 2005; Wenzel et al., 2017). In the latter case, one would predict that activation of host motor axons regenerating into grafts would elicit a wave of excitation in the graft that would be slowly propagated across the graft in the lesion site. Given the benefit of NPC/NSC grafts in improving functional outcomes after SCI in both rodents (Brock et al., 2017; Kadoya et al., 2016; Koffler et al., 2019; Kumamaru et al., 2018; Lu et al., 2012, 2017) and primates (Rosenzweig et al., 2018), it is important to gain an improved understanding regarding which of these mechanisms may be operative.

Thus, to gain insight into the nature of synaptic connectivity and functionality between regenerating host axons and graft neurons in a spinal cord lesion, we optogenetically stimulated host corticospinal axons regenerating into NPC grafts placed into sites of upper lumbar SCI in mice. Graft neurons expressed a genetically encoded calcium indicator, allowing whole-graft imaging of activity in up to nearly one hundred cells simultaneously in spinal cord slices. In addition, we performed live in vivo imaging of calcium activity during host stimulation. Finally, we probed the nature of graft-to-host synaptic connectivity by optogenetically stimulating graft-derived axons innervating host spinal cord neurons caudal to the lesion site that expressed calcium indicators, in slices. We now find that regenerating host corticospinal axons elicit robust and widespread calcium responses in graft neurons, generating correlated activity in clusters of graft neurons in a similar fashion to corticospinal activation of endogenous circuits in intact spinal cords. In turn, stimulation of graft axons extending caudal to the lesion site also activates host spinal neurons in slices. Moreover, in vivo calcium imaging reveals that graft neurons respond to sensory stimuli delivered to the host, including light touch, pinch, and limb movement. These findings show that neural progenitor cell grafts form active synaptic networks within sites of SCI that functionally integrate with host spinal and supraspinal neuron populations, and that resemble physiological patterns of corticospinal projections to the normal spinal cord.

## RESULTS

### Experimental Design

We injected adeno-associated viral vectors (AAV) expressing the red-shifted channelrhodopsin ChrimsonR under the pan-neuronal synapsin promoter (AAV-Syn-ChrimsonR-tdTomato) into the motor cortex in forty-two C57BL/6 mice (Klapoetke et al., 2014); robust expression was subsequently observed in corticospinal axons in the lumbar spinal cord (**Figure 1A-1D, 1F**). Six to eight weeks later, mice underwent T12 dorsal column spinal cord lesions to transect the corticospinal motor projection. In the same session, embryonic day 12 (E12) spinal cord multipotent neural progenitor cells (NPCs) that expressed the calcium indicator GCaMP6f (Chen et al., 2013) in a graft-specific, Cre-dependent manner were grafted directly into the lesion site, as previously described (Adler et al., 2017; Kadoya et al., 2016) (**Figure 1D** and **1E**; **Figure S1**; see Methods). Six to ten weeks post-injury/grafting, when grafts were mature, animals were prepared for either live spinal cord slice (N = 27 animals) or in vivo (N = 15 animals) calcium imaging, as described below. For animals undergoing slice preparation, most of the thoracic and lumbar cord was prepared in the sagittal plane. After imaging 13 animals with graft-specific GCaMP6f expression in slices, it was determined that graft boundaries could faithfully be delineated in a conservative fashion by gross morphology (**Figure S1**). Thus, for later experiments relying on imaging of a larger proportion of graft neurons to analyze activity patterns, we injected non-Cre-dependent GCaMP6f vectors either at the time of grafting or 4 weeks later.

**Figure 1:**
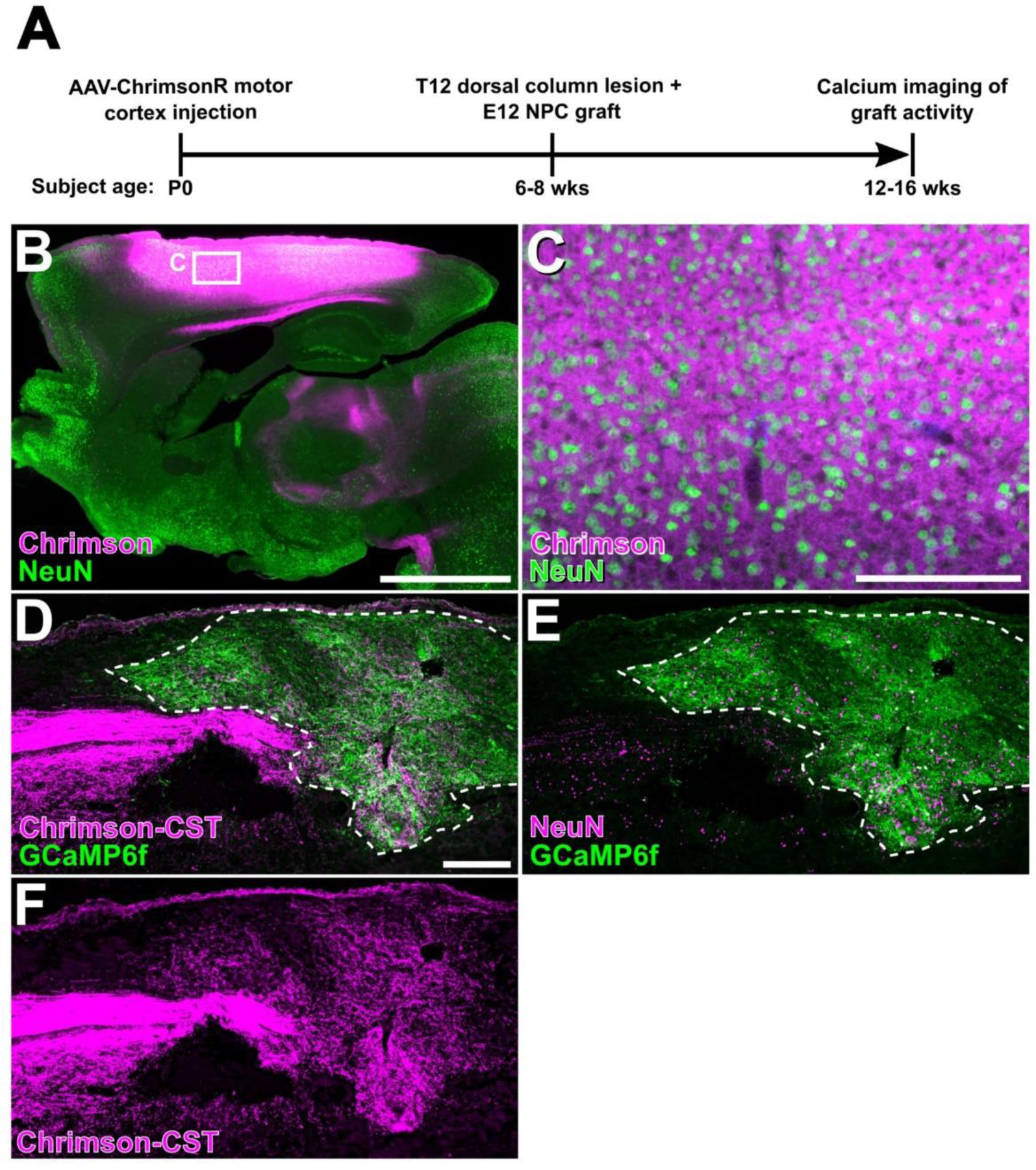
Chrimson-expressing corticospinal axons regenerate robustly into GCaMP6f-expressing neural progenitor cell grafts. (A) Experimental timeline. Animals received cortical injections of AAV-ChrimsonR at postnatal day 0 (P0). T12 dorsal column lesions and acute neural progenitor cell grafting were performed at 6 – 8 weeks of age. Calcium imaging was performed 6 – 8 weeks later. (B) Sagittal section of adult whole brain shows abundant expression of ChrimsonR-tdTomato in cortex following neonatal AAV injections. Neurons labeled with NeuN. Inset in (B) shown at higher magnification in (C). (D) Chrimson-labeled corticospinal tract (CST) axons regenerate extensively into neural progenitor cell grafts expressing GCaMP6f in neurons in a graft-specific, Cre-dependent manner. GCaMP6f (E) and Chrimson (F) in separate channels. NeuN labels neurons in (E). In all images, the dorsal aspect of the slice is at the top and the rostral aspect is to the left. Scale bar, 2 mm (B); 200 µm (C-F).

### Spinal Cord Slices

Using a custom filter set and a standard widefield electrophysiology microscope, corticospinal axons regenerating into neural progenitor grafts were stimulated with a 617 nm LED through the objective lens, and grafts were imaged for neural activity (**Figure S2A**). Regions of interest (ROIs) with neuronal morphology and dynamic fluorescence were manually outlined (**Figure S2B-2C**). The number of detectably active neurons in a field of view was in the range of 10-100 cells. In standard artificial cerebrospinal fluid (aCSF) recording solution, grafts displayed spontaneous activity together with responses to stimulation of corticospinal axons that regenerated into the graft occupying the lesion site (**Figure S2D**). Addition of the potassium channel blocker 4-aminopyridine (4-AP, 100µM; (Blight et al., 1991; Liu et al., 2017; Strupp et al., 2017)), a drug approved for human use, to the aCSF increased detectable spontaneous activity and responses to corticospinal stimulation (**Figure S2E-2G**), and was used in most recordings. Episodes of activity in individual neurons were defined as periods of increasing fluorescence (positive first derivative of the change in fluorescence over time, ΔF/F, after noise subtraction) during fluorescence transients that were significantly above noise based on a dynamic threshold incorporating the individual cell’s calculated noise model and the decay kinetics of the calcium reporter (Romano et al., 2017).

### Spontaneous Activity: Grafts Contain Organized Assemblies of Active Neurons

Graft neurons exhibited spontaneous activity in single cells as well as in clusters of cells that were activated in broad waves of activity at a rate of 2.33 ± 1.10 activations/min (n = 15 clusters in N = 4 animals) (**Figure 2A-2C, Figure S3**). Clusters of neurons that exhibited temporally correlated calcium activity, and that continued to be activated repeatedly over time as the same clusters of temporally associated cells, were termed “assemblies”. These assemblies typically consisted of 5.7 ± 0.5 cells that were usually located close to one another, with a mean distance of 118 ± 27 µm between cells in an assembly (**Figure 2B;** compare to 248 ± 6 µm between cells in random surrogate control assemblies, see below). The PCA-promax method of clustering synchronously active cells (Hendrickson and White, 1964; Peyrache et al., 2010; Romano et al., 2017) demonstrated that most individual neurons within grafts belonged to only one cell assembly as opposed to multiple, different assemblies (only 4.7 ± 3.1% of cells belonged to more than one assembly). Each slice of graft contained roughly 4 separate cell assemblies. This largely independent arrangement of assemblies suggests a degree of compartmentalization in the functional organization of grafts.

**Figure 2:**
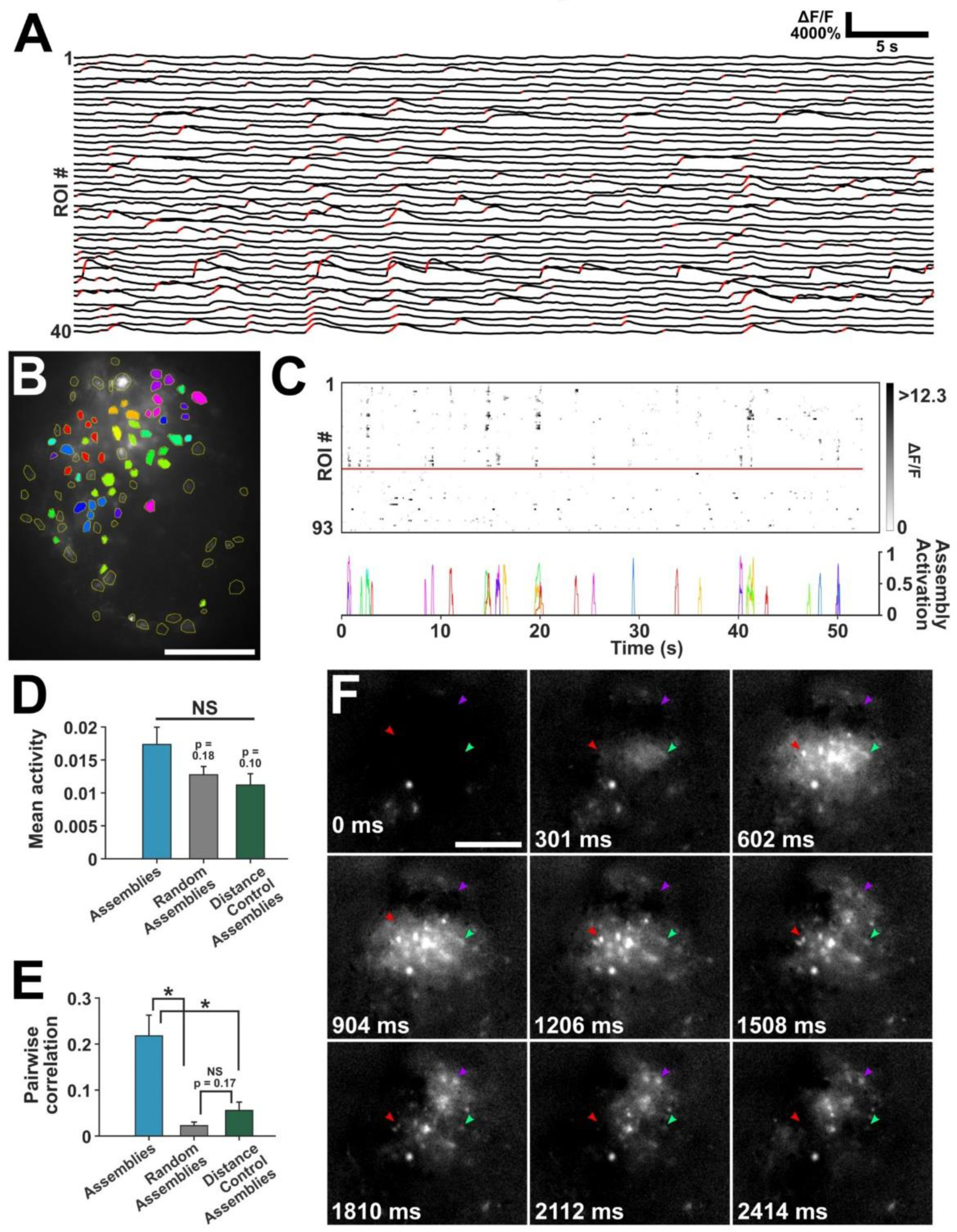
Clusters of graft neurons exhibit spontaneous, correlated activity. (A) Fluorescence traces of 40 graft cells exhibiting spontaneous activity. Red portions of traces indicate periods of significant activity as determined by first derivative rasterization. Individual regions of interest exhibit the onset of calcium activity concurrently, and intermittently repeat this concurrent activation pattern. Moreover, many of these cells are located adjacent to one another, as clusters of cells in grafts (see panel B). Additional traces in Figure S3. (B) Standard deviation projection of the calcium activity run shown in (A) with all regions of interest (ROIs) outlined in yellow; clusters of neurons with similar activity dynamics are indicated by fill color. Many cells exhibiting concurrent onset of calcium activity are spatially adjacent to one another; we refer to these clusters as “assemblies.” (C) First derivative raster plot (top) of traces from (A), with cells grouped into the assemblies shown in (B). Cells above red line belonged to assemblies. Significant activations of color-coded assemblies are plotted below. (D) The mean activity of cells belonging to assemblies was not significantly different from that in control, random assemblies or surrogate distance-control assemblies (see text; N = 4 animals per group analysis). (E) Cells in assemblies showed significantly more pairwise correlation than those of surrogate assemblies (N = 4 animals). (F) Series of serial imaging frames (ΔF/F) showing temporal rise of calcium activity in separate assemblies in the same imaging run as in (A-C). Generally, peak calcium activity is reached over ∼500-1000 msec. Arrowheads indicate same cells within three separate assemblies over sequential frames. The colors of the arrowheads indicate the assemblies in (B) to which the cells belong. In all images, the dorsal aspect of the slice is at the top and the rostral aspect is at left. Scale bar, 200 µm (B, F). Data presented as mean ±SEM with significance determined by Welch’s t-test (*p<0.05; NS, not significant).

To confirm the validity of cell assemblies that we identified based on patterns of concurrent spontaneous activity, we generated control “surrogate” assemblies from the total pool of regions of interest (ROIs) in each slice (Romano et al., 2017) as follows: at the beginning of the analysis pipeline, neurons were numbered by their location in the field of view from left to right. An assembly of neurons with temporally correlated activity could therefore be described by a list of numbers of the neurons belonging to that assembly. Surrogate assemblies were then generated by preserving these lists but shuffling the original numbering of the total pool of neurons. Thus, sets of surrogate assemblies contained the same quantity of neurons in each assembly as the experimentally derived set. These surrogate control assemblies were either: a) completely random groupings (“random assemblies”), or b) groupings that maintained the same distances between cells as the experimentally identified assemblies, but comprised of different sets of cells than the experimentally defined assemblies (“distance control assemblies”). One hundred random surrogate control and 100 distance surrogate control assemblies were generated for each experimentally identified assembly, and these control data sets were compared to experimentally identified assemblies. The total pool of experimentally identified assemblies across animals had similar levels of mean activity per cell compared to surrogates, but much higher pairwise activity correlation between cells, indicating that the clusters were not merely groups of cells with increased activity or close proximity but assemblies of cells grouped either by direct connectivity or similar pre-synaptic inputs (Carrillo-Reid et al., 2016; Hebb, 1949; Romano et al., 2015; Yuste, 2015) (**Figure 2D-2E**).

### Temporal Spread of Activity Within Graft Neuronal Assemblies

We next examined the temporal dynamics of spontaneous activity in grafts within individual neuronal assemblies by examining sequential calcium imaging in 300 msec (10 imaging frame; **Figure 2F**) and 30 msec (single imaging frame, **Figure S4**) time epochs. Calcium activity appeared to arise simultaneously (within the temporal resolution of calcium imaging) within single neuronal assemblies, reaching peak calcium activity within the cluster over a time period of 500 – 1000 msec (**Figures 2F** and **S4**). The coordinated rise in calcium activity among neurons belonging to assemblies in the grafts suggests that they are highly interconnected. This pattern of coordinated rise in calcium activity within a cell assembly is similar to the pattern observed in the intact spinal cord, below.

### Evoked Activity: Corticospinal Inputs Stimulate Network Activity in Graft Neuronal Assemblies

Next, we sought to identify patterns of synaptic organization in grafts following stimulation of host corticospinal axons regenerating into grafts. Graft neurons displayed robust calcium responses to corticospinal optogenetic stimulation within hundreds of milliseconds of stimulus onset (**Figure 3A-3D**). At the low magnification used for population calcium imaging, subthreshold fluctuations in membrane potential are not generally detectable above noise, whereas action potentials can generally be resolved with the calcium indicator GCaMP6f (Emiliani et al., 2015; Lin and Schnitzer, 2016). We therefore interpret the fast-rising and slow-decaying fluorescence transients observed in response to corticospinal stimulation to be the result of one or more action potentials firing in graft neurons. The fluorescence traces in responding cells began to rise immediately following stimulus onset and reached their temporal midpoint to peak fluorescence less than halfway through the 500 msec stimulus (**Figure 3E**), indicating that the responses saturate in less than half a second. These responses were mediated through excitatory transmission, as indicated by their disappearance in the presence of the glutamatergic blocker 6,7-dinitroquinoxaline-2,3-dione (DNQX) (**Figure 3E-3F**). The magnitude of response strength (average ΔF/F over 1 sec following stimulus onset) and percent of responding cells (ΔF/F ≥ 100%) increased with increasing power of the LED stimulation (**Figure 3G-3H**), demonstrating that graft responses encode stimulus strength. When AAV-Chrimson was not injected into the cortex, graft cells did not respond to LED stimulation in the absence of opsins (N = 6 animals; **Figure S2H**), indicating that graft responses to stimulation are not artifacts of direct photoactivation by the light stimulus.

**Figure 3:**
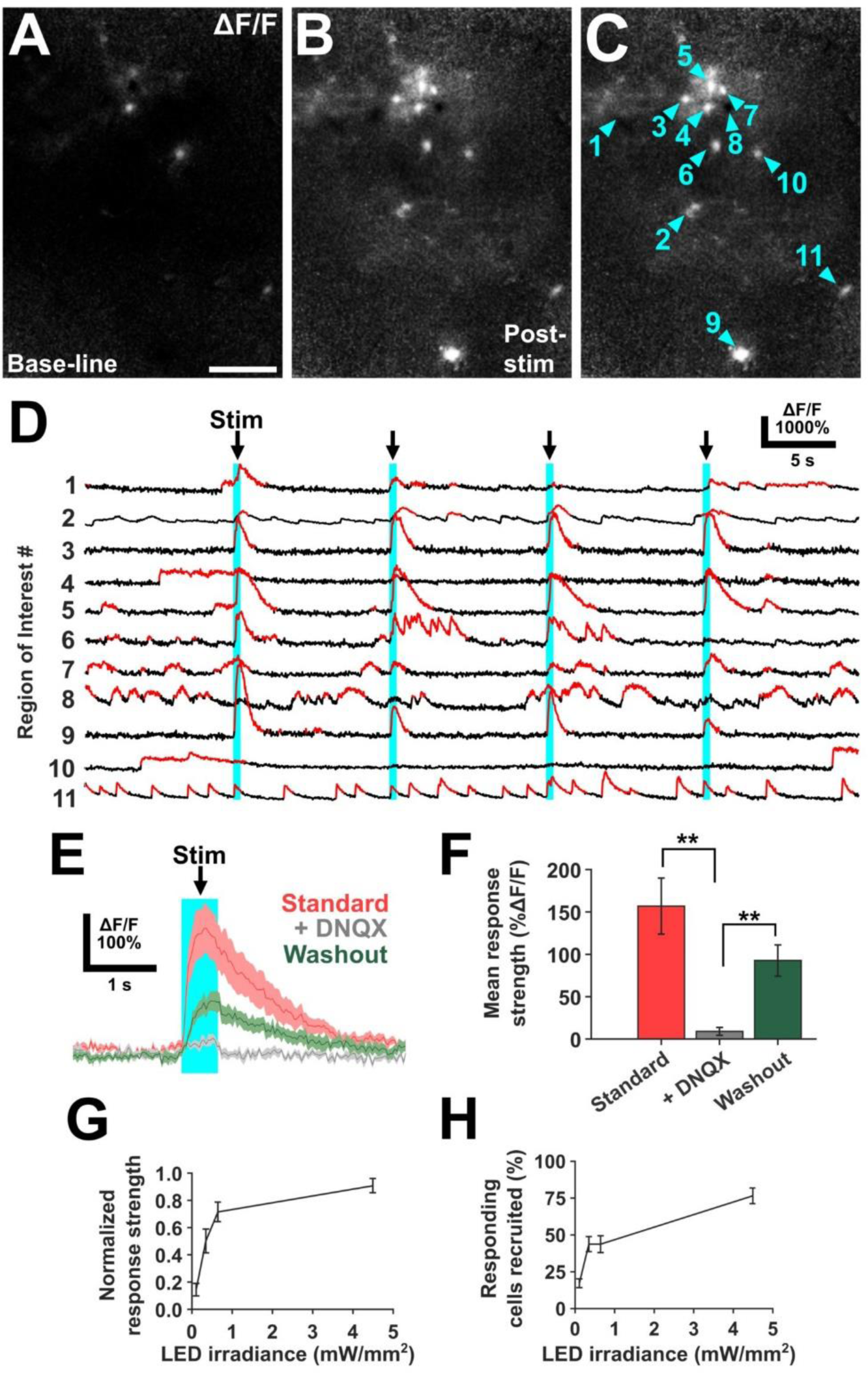
Graft neurons are activated by optogenetic stimulation of regenerating corticospinal axons. (A) One second average of ΔF/F video prior to stimulation onset shows several spontaneously active graft neurons. (B-C) The same field as (A), immediately following stimulation onset. GCaMP6f fluorescence is activated in several graft neurons in response to stimulation of corticospinal axons that have regenerated into the graft (cyan numbered arrows in (C)). (D) Sample traces from cells labeled in (C) during a 50 sec period over which four corticospinal stimuli (vertical cyan bars) are delivered. Stimuli consisted of 500 msec-long 20 Hz trains of 10 msec pulses of 617 nm light. Irradiance at the slice was 4.49 mW/mm^2^. Significant fluorescence transients are highlighted in red. Cells display varying response strengths as well as spontaneous activity. Notably, clusters of neurons within grafts repeatedly responded to corticospinal stimulation (see also Fig. 4). (E) Mean response traces with ±SEM shade of n = 7 cells in one slice to 12 trials in standard recording conditions, + 100 µM 6,7-dinitroquinoxaline-2,3-dione (DNQX), and after DNQX washout. Responses were abolished by the AMPA and kainite receptor antagonist DNQX, indicating the presence of excitatory synaptic connections from corticospinal axons onto graft neurons. (F) Mean response strengths from (E), quantified over one second following stimulus onset. (G) Mean response strength quantified as in (F) in response to increasing 617 nm LED irradiance. Responses in n = 16 cells in one slice increased in strength with increasing irradiance and plateaued at higher irradiance. (H) The percentage of cells activated (activation threshold = 100% ΔF/F) by corticospinal stimulation also increased with increasing 617 nm LED irradiance. In all images, the dorsal aspect of the slice is at the top and the rostral aspect is at left. Scale bar, (A-C), 100 µm. Two-group comparisons were tested with Welch’s t-test (**p<0.01; NS, not significant). Data are represented as mean ±SEM.

Cells throughout grafts responded to corticospinal stimulation, even at the most caudal aspect, furthest from the point at which the main tract of corticospinal axons enters the graft (**Figure S5**). Such caudal responses began immediately following stimulus onset, not exhibiting a delay in latency compared to more rostral responses. While the density of ChrimsonR-tdTomato-labeled axons varied throughout different regions of grafts, as previously observed (Dulin et al., 2018; Kumamaru et al., 2019), response strength to corticospinal stimulation in a given cell was uncorrelated with the brightness of tdTomato signal in the corresponding region of interest (n = 68 cells in N = 3 animals; response strength and tdTomato brightness normalized to their respective maximum in each imaging field; **Figure S5D-S5E**). Thus, graft cells are activated by corticospinal stimulation in all regions of grafts, including the caudal regions of grafts that contains the preponderance of graft neurons that extend axons into the host spinal cord below the injury (Lu et al., 2019).

### Corticospinal Axons Activate the Same Neuronal Assemblies that Were Identified Based upon Spontaneous Activity

Optogenetic stimulation of corticospinal axons within grafts activated *the same graft neuronal assemblies* that were identified based on analysis of spontaneous activity in grafts. Indeed, 22.5 ± 13.2% of clusters that displayed spontaneous activation in a 50 sec imaging run also responded to corticospinal optogenetic stimulation (n = 18 clusters in N = 4 animals, **Figure 4A-4B**). The overall topography of activation was similar with each stimulation trial (**Figure 4C-4H**), although some individual cells and clusters exhibited responses in only a portion of trials. Assembly activation by corticospinal stimulation occurred simultaneously in all responding cells of an individual assembly, and stereotyped sequential activation within an individual assembly was not observed.

**Figure 4:**
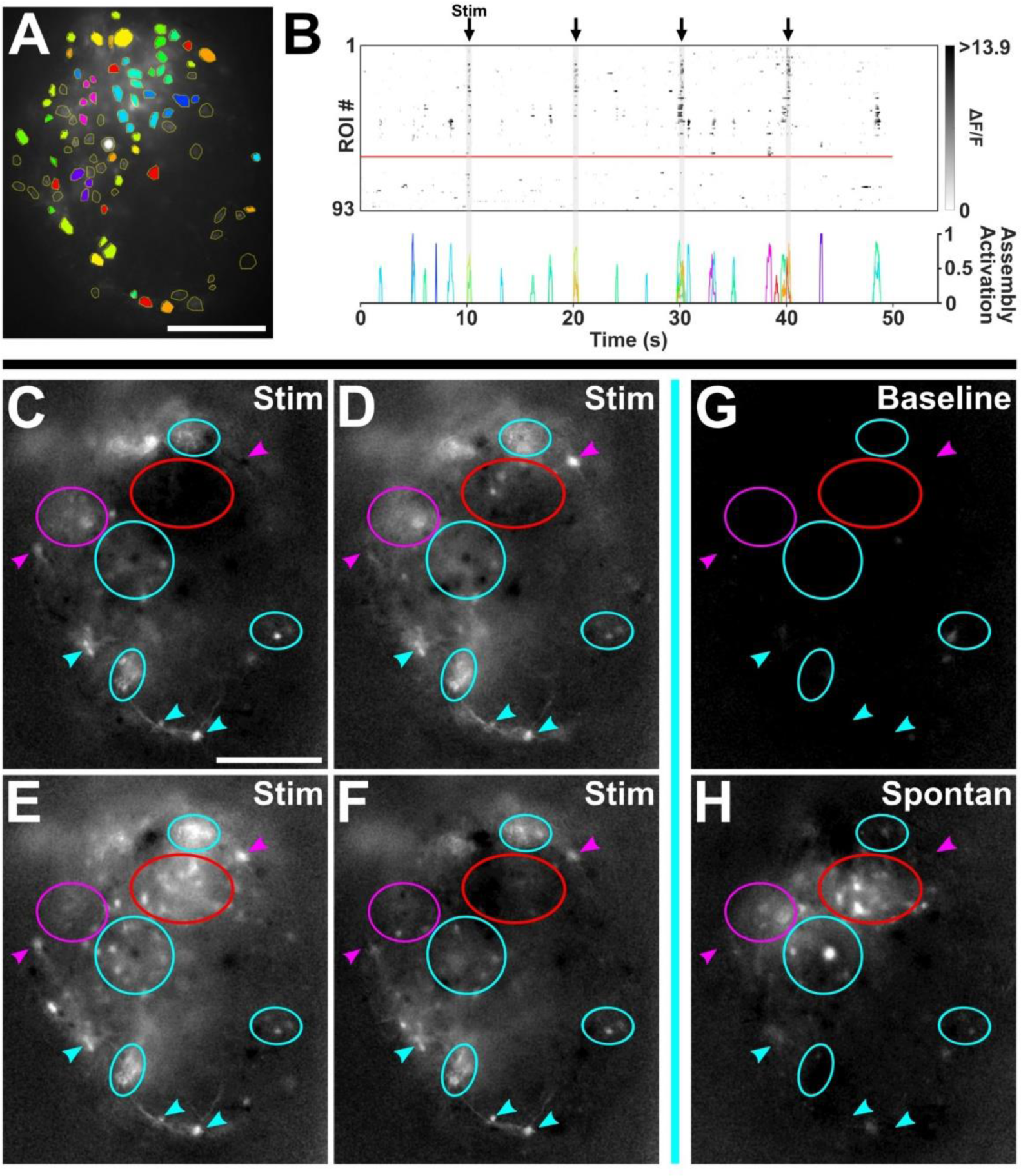
Spontaneously active graft neuron assemblies are activated by corticospinal stimulation. A) Same graft as in (Figure 2B) with assemblies extracted from a run with optogenetic corticospinal stimulation. (B) As in (Figure 2C), during a run with four trials of 500 msec, 20 Hz corticospinal stimulation. (C-F) One second, serial 33-frame average ΔF/F images of graft responses to four consecutive corticospinal stimulation trials show graft regions (ellipsoids) and individual neurons (arrowheads) that respond consistently (cyan) or inconsistently (magenta) to stimulation. Red circle indicates a region that was spontaneously active but did not respond to corticospinal stimulation (region was already activated prior to stimulus in (E)). (G) One second average from the same field of view as (C-F) during a period of low activity in an imaging run with no stimuli given. (H) The same field during a period of spontaneous activation of the regions in the red and magenta ellipsoids. In all images, the dorsal aspect of the slice is at the top and the rostral aspect is at left. Scale bar, 200 µm (A,C-H).

### Whole Grafts Do Not Exhibit Slow Spreading of Activity

It is possible that host inputs regenerating into grafts could elicit a slow, spreading wave of sustained depolarization through grafts (Ayata and Lauritzen, 2015). However, we observed no evidence for the existence of this mechanism: imaging grafted spinal cord slices following corticospinal activation in a total of 14 animals with responding grafts, we never observed a “spreading wave” of calcium activation. Rather, corticospinal activation evoked simultaneous activity in discrete neuronal assemblies within grafts (**Figure 4**), indicating that grafts organize into functional synaptic networks that repeatedly and stereotypically respond to activation of host inputs. Such activation could relay excitation to host spinal circuits below the injury, transmitted by the tens of thousands of axons that emerge from grafts into the caudal host (Lu et al., 2012, 2014, 2017; Rosenzweig et al., 2018) (**Figure 5A-5B**).

**Figure 5:**
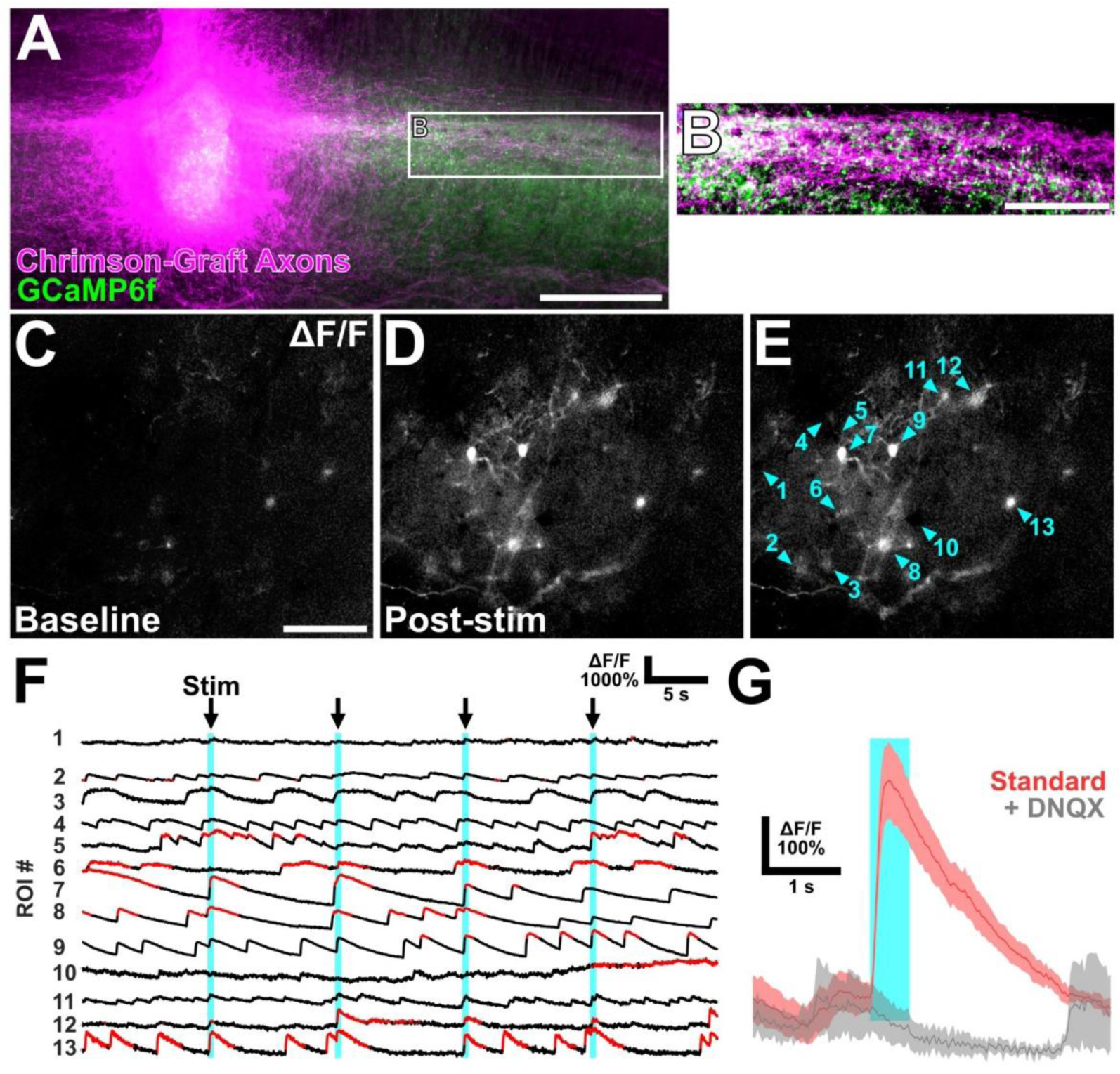
Host neurons are activated by optogenetic stimulation of axons extending from grafts. (A) Graft that expresses Chrimson extends axons into host spinal cord rostral and caudal to the lesion site. (B) Inset from (A) showing graft axon innervation of regions of GCaMP6f-expressing host spinal cord neurons caudal to the lesion. (C) One second average ΔF/F image of host spinal cord neurons prior to stimulus onset. (D-E) In the same field as (C), one second average following onset of 500 msec optogenetic stimulation of graft axons (cyan numbered arrows in (E) label analyzed cells). (F) Individual cell fluorescence traces of the cells in (E) show abundant spontaneous activity as well as consistent responses in some cells to graft axon stimulation. (G) Mean ±SEM traces of n = 5 responding cells from one slice before and after wash-in of 100 µM DNQX, showing that grafts form excitatory synaptic connections with host neurons below the lesion. In all images, the dorsal aspect of the slice is at the top and the rostral aspect is at left. Scale bars, 300 µm (A); 150 µm (B); 100 µm (C-E).

### Intact Spinal Cords are Also Organized in Neuronal Assemblies

To assess whether graft responses to host inputs resemble patterns evoked by corticospinal inputs to the intact spinal cord, we also examined (in slices) responses of spinal cord neurons to optogenetic stimulation of corticospinal neurons innervating the intact spinal cord. Four animals underwent post-natal day 1 injections of AAV-Syn-ChrimsonR-tdTomato into the primary motor cortex, followed by injections of AAV vectors expressing GCaMP6f into the T12 spinal cord at 2 months of age. During acute slice imaging one month later, spinal cord neurons of the medial and dorsal gray matter displayed spontaneous activity in both individual neurons and in clusters of neurons with correlated activity; as in neural progenitor grafts, cells exhibiting correlated activity were spatially close to one another (**Figure S6** and **S7C**) and were activated repeatedly over time (**Figure S7C**). When the corticospinal tract was optogenetically stimulated, roughly a quarter of the same assemblies of neurons in the spinal cord exhibited calcium activity (**Figure S7D**). Like responses observed in neural progenitor grafts, frame-by-frame analysis of the temporal profile of spinal cord response to corticospinal activation revealed simultaneous rather than sequential activation of neuronal assemblies in the intact spinal cord.

### Graft-to-Host Activity

Having observed this activation of assemblies of cells throughout grafts following stimulation of regenerating corticospinal axons, we next sought to determine whether grafts could in turn activate *host* spinal neurons caudal to the injury site. We mixed AAV-Syn-FLEX-ChrimsonR-tdTomato vectors with Syn1-Cre neural progenitor cells just prior to grafting acutely into T12 dorsal column lesions. This yielded graft-specific expression of ChrimsonR-tdTomato, which was readily detectable in *axons* extending out from the graft borders and into the distal host spinal cord (**Figure 5A-5B**). Three to four weeks after grafting, AAV9-Syn-GCaMP6f was injected 1.2 mm caudal to grafts into host central gray matter (**Figure 5A-5B**). When imaged 6-8 weeks after grafting, a total of 15 host neurons were detected among slices from two animals that responded to whole-field stimulation of graft axons (**Figure 5C-5F**). Responding cells were located in the medial and dorsal gray matter. Host responses to graft axon stimulation were eliminated after DNQX administration, indicating that host responses to graft axon stimulation were mediated through excitatory synaptic transmission (**Figure 5G**). The latency of these responses was less than 200 msec, showing that, as in grafts, responses peaked before the end of the 500 msec light stimulus (**Figure 5G**). Thus, neural progenitor cell grafts are capable of activating host neurons below spinal cord lesions.

### In Vivo Calcium Imaging Shows Functional Integration of Host with Graft

To examine activation of graft neurons by behavioral stimulation of the host, we turned to in vivo imaging of graft activity (Farrar et al., 2012; Sekiguchi et al., 2016). Animals underwent T12 dorsal column spinal cord lesions followed by placement of E12 neural progenitor cell grafts expressing GCaMP6f into the lesion site. Six to ten weeks later, spinal stabilization hardware and imaging windows were implanted over the dorsal surface of grafts, as previously described (Farrar et al., 2012; Sekiguchi et al., 2016). Grafts were imaged one hour after window implantation in anesthetized animals. With a detectable depth of imaging of approximately 100 µm into grafts, we observed spontaneous activity of graft neuron cell bodies (**Figure 6A-6C**) and processes (**Figure 6D-6G**). Up to ten regions of interest exhibiting responses to host simulation were visible in a single field of view. In anesthetized animals, when the host was stimulated by hindpaw pinch (**Figure 6D-6E**), light touch of the lower back (**Figure 6F-6G**), or passive movement of the hindlimb (**Figure 6H**), individual graft neuronal responses were detected in four out of seven animals with detectable graft activity. These responses could be consistently reproduced over repeated stimuli, some exhibiting sensitization of response strength to the stimulus and others exhibiting desensitization or no change in response strength (**Figure 6E, 6G-6H**). Cells typically responded to a single stimulus type, though two neighboring ROIs responded to both hindlimb movement and light touch of the lower back in one animal (**Figure 6F-H**). Thus, host sensory systems functionally connect to grafts and elicit synaptic responses.

**Figure 6:**
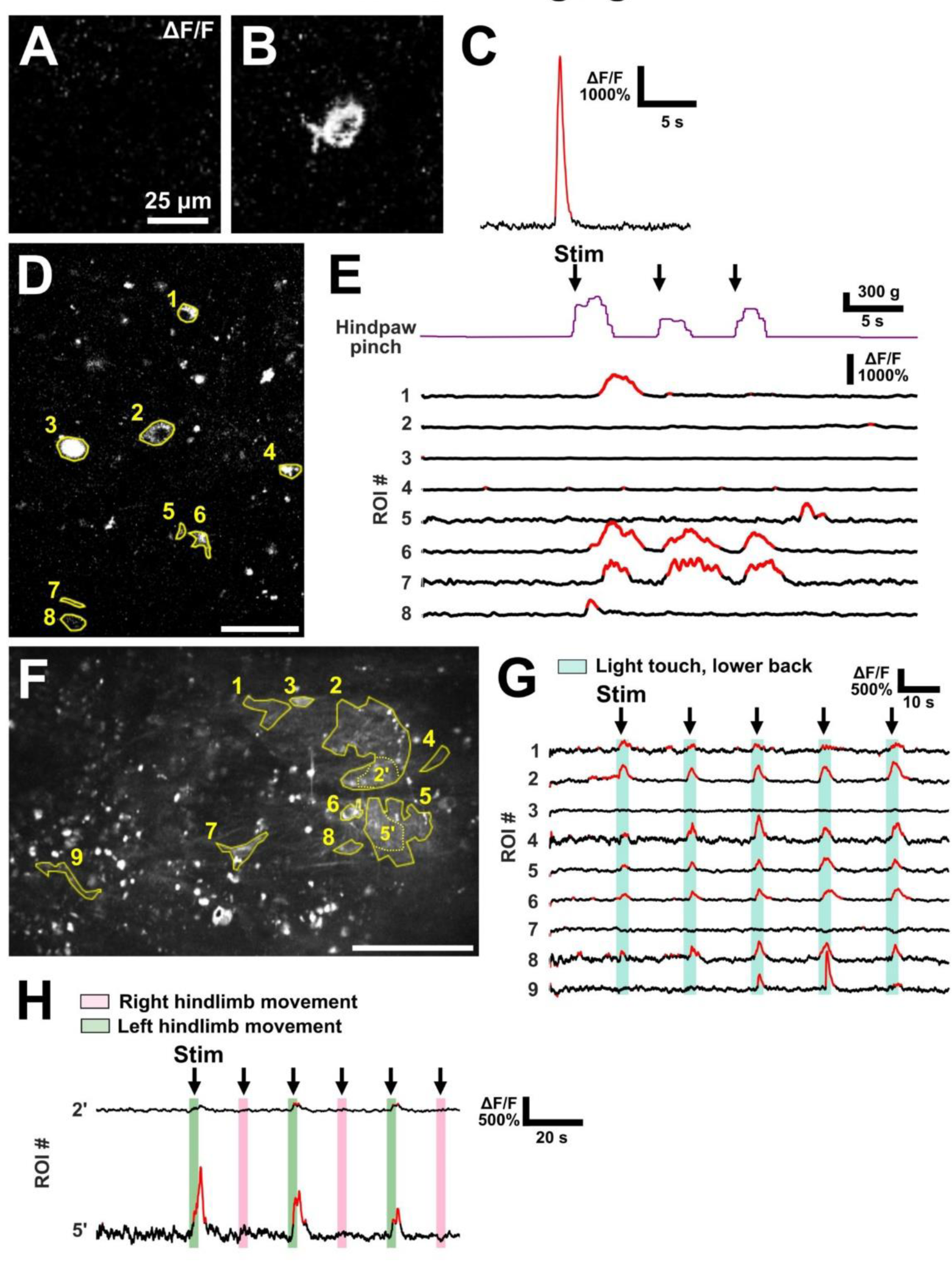
Graft neurons respond to sensory stimuli in vivo. (A-B) Baseline average ΔF/F image of a graft neuron over a one second period just prior to (A) and one second following onset of a spontaneous calcium transient (B) in a single graft neuron with graft-specific GCaMP6f expression in an anesthetized animal. (C) Fluorescence trace of the calcium transient shown in (A-B). (D) Maximum projection image of in vivo calcium activity during a period of three hindpaw pinches. ROIs (yellow outlines) include both neuronal soma and processes. (E) Fluorescence traces of the activity in the ROIs shown in (D) during hindpaw pinch (purple trace). (F) Standard deviation projection of viral GCaMP6f expression in vivo during light touch. ROIs are highlighted in yellow. (G) Several ROIs respond consistently to light touch. (H) Two cropped ROIs from the same field of view as (E) respond consistently to movement of the hindlimb through its range of motion by the experimenter. These ROIs, which may be part of a single neuron or neuron cluster, were on the left side of the graft and responded to left hindlimb movement (green bars) but not right hindlimb movement (pink bars). Scale bars, (A-B), 25 µm; (D), 30 µm; (F), 100 µm.

## DISCUSSION

We report now for the first time that host axons regenerating into neural progenitor cell grafts placed into sites of SCI activate responses from highly correlated assemblies of graft neurons, in addition to individual graft neurons. A previous elegant two-photon study demonstrated that host axons regenerating into stem cell grafts in the cortex could elicit stimulus-specific responses from individual neurons (Falkner et al., 2016), but up to the present no studies have been performed that examine the nature of broader graft functional synaptic network properties. The activation of neuronal assemblies within grafts that we now identify resembles responses observed after optogenetic stimulation of corticospinal axons in the intact spinal cord. Host corticospinal axons regenerating into grafts do *not* generate slow waves of excitation that spread through grafts, indicating that graft responses are more similar to those seen in healthy neuronal networks rather than dysfunctional disease states. Developing neural circuits exhibit rhythmic waves of spontaneous activity that are synchronous in a large proportion of cells, and these patterns generally cease by the end of the first week of post-natal development (Blankenship and Feller, 2009; Garaschuk et al., 2000); in contrast, we observed more mature, discrete cell cluster activity from neural progenitor cell grafts at the equivalent of time points of five weeks and later postnatally. Considering the highly anomalous milieu that we are studying, i.e., “developing” neural tissue placed in an *adult* spinal cord lesion site, it is remarkable that spontaneous graft activity and the patterns of synaptic connectivity recruited by regenerating host axons into grafts generally resemble patterns of the intact spinal cord. Moreover, grafts respond to *behavioral* stimulation of the host in a similar fashion to dorsal horn neurons in intact spinal cord (Johannssen and Helmchen, 2013; Sekiguchi et al., 2016).

Graft neurons respond to corticospinal input as quickly as is detectable with calcium imaging methods. The latency of the temporal midpoint to peak response amplitude was 100-300 ms, and cell fluorescence began to increase within one to two imaging frames (about 30-60 ms) of stimulus onset in cells throughout grafts. Similar response latencies were obtained for host neurons below the lesion to graft axon stimulation. While the underlying electrical activity which gives rise to GCaMP6f fluorescence spikes occurs on a much shorter timescale (Chen et al., 2013), even an extremely conservative fluorescence-based estimation would place the time for a full relay across the lesion at no more than several hundred milliseconds.

In the dorsal column lesion model employed in this study, regenerating corticospinal axons were present throughout the rostro-caudal length of the graft, and corticospinal stimulation activated neurons throughout the graft, including caudal graft regions where fewer corticospinal axons were observed. Similarly, response strength did not depend on local density of Chrimson-expressing fibers. These widespread responses to light stimulation could be due to the dendritic processes of graft neurons extending into areas of denser Chrimson-expressing axon innervation or through polysynaptic connections with other graft cells receiving direct corticospinal input. Among all spontaneously active neuron assemblies in grafts, approximately one-fourth responded to corticospinal stimulation. This limited responsiveness could reflect an incomplete extent of axonal regeneration, but it could also indicate that regenerating corticospinal axons may elicit responses from subdomains of grafts containing appropriate corticospinal target neurons; such subdomains spontaneously assemble in grafts of neural progenitor cells placed in sites of SCI (Dulin et al., 2018; Kumamaru et al., 2019).

Previous studies have examined graft-host connectivity and relay formation by gross extracellular stimulation and recording on either side of a graft (Koffler et al., 2019; Lu et al., 2012), but even when guided stereotactically, this method is ambiguous with regard to the exact cell populations being tested due to potential spread of current through tissue. It is also possible that stimulation of the spinal cord at one level may cause muscle twitches and associated body movement that could result in reflex responses in the spinal cord on the distal side of the lesion, potentially confounding results that indicate the presence of a neuronal relay. The optogenetic approach taken here shows unequivocal host-to-graft and graft-to-host functional connectivity in defined neuron populations, and will prove useful in further dissecting the role of distinct host inputs and graft neuron subtypes in relay action.

## CONCLUSION

Findings of this study indicate that host axons regenerating into neural progenitor cell grafts recapitulate features of neuronal activation of the intact spinal cord. For example, grafts exhibit activation of localized neuronal clusters or assemblies. Grafts also exhibit synaptic transmission of activity from one “spinal cord” region (the graft) to another (the caudal spinal cord); this resembles the polysynaptic and interneuronal features of corticospinal inputs to motor neurons via motor synergy encoder neurons of the intact spinal cord (Alstermark and Isa, 2012; Azim et al., 2014; Levine et al., 2014; Ueno et al., 2018). In vivo calcium imaging shows the functional integration of host inputs to grafts. These findings lend further support to the potential feasibility of transplanting neural stem cells to treat spinal cord injury.

## METHODS

### Animals

Young adult wild-type C57BL/6J [(The Jackson Laboratory, stock number: 000664)] mice (15-25 g, n = 56) were subjects of this study. To generate E12 embryos for neural progenitor cell harvesting, Syn1-Cre [B6.Cg-Tg(Syn1-Cre)671Jxm/J (The Jackson Laboratory, stock number: 003966)(Zhu et al., 2001)] males were bred to either wild-type C57BL/6J or Ai95D females [B6;129S-Gt(ROSA)26Sortm95.1(CAG-GCaMP6f)Hze/J (The Jackson Laboratory, stock number: 024105)(Madisen et al., 2015)]. All procedures were carried out in accordance with NIH guidelines for laboratory animal care and safety and were approved by the Veterans Administration San Diego Healthcare System IACUC. Animals had free access to food and water throughout the study. Surgeries were performed under deep anesthesia using a combination (12.5 ml/kg) of ketamine (2.5 mg/ml), xylazine (0.13 g/ml), and acepromazine (0.025 mg/ml); and 1% isoflurane.

### Chrimson Virus Injection

Neonatal pups (P0-P1) were anesthetized via hypothermia on a wet towel on ice and placed on an ice-cold stereotax. The scalp was rubbed with 70% ethanol before and after the injection procedure. Pups were injected with 0.5 µL of AAV1-Syn-ChrimsonR-tdTomato (1 × 10^12^ vg/mL) in each hemisphere of the motor cortex using a 34 gauge, 0.375” needle with a 12° tip in a Hamilton syringe attached to syringe pump (NanoJet, Chemyx Inc). The narrow gauge and sharp tip of the needle allowed it to penetrate directly into the cortex through the scalp and skull. An injection rate of 0.75 µL/min was used, and 30 seconds were allowed before removing the needle after the injection completed. Pups recovered on a heat pad until pink and wriggling before being rubbed with bedding and feces from their home cage and returned to the mother.

### Cell Preparation

Neural progenitor cells were prepared for grafting from embryonic day 12.5 (E12.5) embryos following a procedure based on previously described methods (Adler et al., 2017). Briefly, embryonic spinal cords were dissected in ice-cold Hank’s Balanced Saline Solution (HBSS), dissociated with 0.05% Trypsin and re-suspended in ice-cold Dulbecco’s Phosphate Buffered Saline (DPBS). Cell viability was assessed by trypan blue exclusion, and aliquots for grafting were prepared at a density of 250,000 cells/µL. In cases where AAV vectors were mixed with graft cells just prior to grafting, virus was added to a final titer of 1.17 × 10^12^ vg/mL + 1.65 × 10^12^vg/mL (AAV1-+ AAV9-CAG-FLEX-GCaMP6f), 8.3 × 10^11^vg/mL (AAV9-Syn-GCaMP6f), or 4.2 × 10^12^ vg/mL (AAV1-Syn-FLEX-ChrimsonR-tdTomato) in the cell suspension.

### Spinal Cord Lesion and Grafting

Dorsal column lesions and acute grafting were performed based on methods previously described (Adler et al., 2017). Briefly, the T12 lamina was removed and a wire knife (McHugh Milieux) was inserted stereotactically to a depth at which the lowest point of the 1 mm-wide extended knife was 1.1 mm below the dorsal surface of the dura mater. Once at this depth, the knife was extended, raised 1.1 mm, and held at this position while the remaining white matter on top of the knife was crushed with a bent insulin syringe needle. The knife was then lowered to its original depth, retracted, and removed from the spinal cord. Neural progenitor cells were immediately grafted using pulled glass micropipettes and a Picospritzer III (Parker Hannifin). Cells were injected directly into the lesion center in a volume of 2 µL DPBS.

### GCaMP6f Virus Injection

AAV9-Syn-GCaMP6f (0.5 µL/site, 1 × 10^12^ vg/mL) was injected either directly into the graft center or into the spinal cord gray matter 0.3 mm from midline (bilaterally) over a depth of 1.2 to 0.6 mm with a pulled glass micropipette and Picospritzer.

### Live Spinal Cord Slice Preparation

Acute spinal cord slices were prepared following a protocol modified from a previously described procedure (Husch et al., 2011). Briefly, animals were placed under deep anesthesia with (28 mL/kg) ketamine (2.5 mg/ml), xylazine (0.13 g/ml), and acepromazine (0.025 mg/ml) and moved to a bed of ice. A block of the spinal column containing the thoracic and lumbar segments of the spinal cord was quickly removed and placed into an icy slurry of oxygenated (95% O_2_-5% CO_2_) artificial cerebrospinal fluid formulated specifically for dissecting spinal cord (DaCSF) containing (in mM): 2 kynurenic acid, 191 sucrose, 0.75 K-gluconate, 1.25 KH_2_P0_4_, 26 choline bicarbonate 80% solution, 20 dextrose, 1 (+)-sodium L-ascorbate, 5 ethyl pyruvate, 3 myo-inositol, 4 MgSO_4_, and 1 CaCl_2_ (∼310 mosmol/kgH_2_O, pH 7.35). The animal was then decapitated with scissors. After transferring the spinal block to a dissecting dish with an icy slurry of DaCSF with constant oxygenation, the spinal cord was dissected from the column, with special care taken to cut any scar tissue stuck between the cord and the laminae and attached tissue without pulling on the graft. Due to the presence of scar tissue over the graft site, we did not remove the dura mater prior to slicing. The spinal cord was then transferred to a custom-built chamber filled with 34°C low melting point agarose (0.03 g/ml in DaCSF; A0701; Sigma), and the agarose was allowed to gel on ice with the cord positioned with the lateral aspect facing up. Once gelling completed, the chamber was transferred to an icy DaCSF slurry and the agarose block was trimmed and transferred to the vibratome buffer tray containing continuously oxygenated ice-cold DaCSF. Slices (330 µm thick) were then prepared in the sagittal plane using a Leica VT1000S vibratome and placed with a fine paint brush into a recovery chamber where they were submerged in 34°C oxygenated DaCSF for 30 min. Slices were then transferred to 34°C oxygenated recording buffer (RaCSF) containing (in mM): 121 NaCl, 3 KCl, 1.25 NaH_2_PO_4_, 24 NaHCO_3_, 1.1 MgCl_2_, 2.2 CaCl_2_, 15 dextrose, 1 (+)-sodium L-ascorbate, 5 ethyl pyruvate, and 3 myo-inositol (∼310 mosmol/kgH_2_O, pH 7.35) for 30 min. The RaCSF recording chamber was then transferred to room temperature, where the slices remained submerged under oxygenation until transfer to the recording chamber.

### Slice Imaging

Spinal cord slices were transferred to a submersion type chamber and perfused with oxygenated RaCSF (with or without 100 µM 4-AP) at 34°C. Ten minutes of recovery time was allowed in the recording chamber prior to imaging. Grafts were identified by fluorescence and/or morphology under infrared differential interference contrast (IR-DIC). Corticospinal axon regeneration or graft axon extension were briefly assessed by ChrimsonR-tdTomato fluorescence under a mercury lamp (U-RFL-T, Olympus), after which only low power LED excitation of GCaMP6f was used for imaging to avoid unintentional activation of Chrimson molecules. The slice imaging rig consisted of (in brief) a widefield fluorescence microscope (BX51WI, Olympus), a 10X NA 0.3 water immersion objective (UMPLFLN10X/W, Olympus), blue (470 nm, 750 mW) and orange (617 nm, 650 mW) mounted LEDs (Thorlabs), two 1200 mA LED drivers (Thorlabs), two NA 0.6 aspheric condenser lenses (ACL2520U-A, Thorlabs), 470 nm and 620 nm excitation filters (ET470/40X and AT620/50X, Chroma), a longpass dichroic mirror with 490 nm cutoff (DMLP490R, Thorlabs), a custom filter cube containing a 530 nm dichroic bandpass and 525 nm emission filter (zt530/55dcbp and ET525/50m, Chroma), and a Rolera XR Fast 1394 or Retiga Electro CCD camera (QImaging). Resolution was 348 × 260 pixels with the Rolera XR and 688 × 512 pixels with the Retiga Electro, both with 2 × 2 binning. Blue LED excitation and orange LED stimulation were controlled with a MultiClamp 700b patch-clamp amplifier via a Digidata 1440A digitizer using Clampex software (all from Molecular Devices). Still and video images (33 frames per second) were acquired with Micro-Manager software (Edelstein et al., 2014). Blue excitation irradiance at the slice was kept low at 0.093 mW/mm^2^ to avoid off-target Chrimson stimulation, and orange stimulation irradiance was 4.49 mW/mm^2^ at maximum current.

### In Vivo Imaging Hardware Implantation

The week prior to imaging, spinal stabilization hardware was implanted as previously described (Farrar et al., 2012). Briefly two magnetic stainless steel bars were placed against the lateral aspect of the spine, centered around the graft site at the T12 spinal level. A stainless steel plate with an opening over the spinal cord was screwed onto the bars and cyanoacrylate glue and dental cement were used to secure the hardware and attach the skin to it to seal the tissue from the environment. On the day of imaging, glue was removed from the surface of the spinal cord. Typically, the dura mater was resected due to scar tissue from the injury preventing clear imaging. A No. 0 glass coverslip was placed over the exposed cord with the graft in the center. Light pressure was applied to the coverslip while it was glued in place to maintain a flush interface with the spinal cord, and the glue was allowed to dry for one hour prior to imaging.

### In Vivo Imaging

All in vivo imaging experiments were acute, terminal procedures. Mice were placed under gas anesthesia with 1.75% isoflurane and restrained at the spine with two rods attached to the spinal plate. They were then placed under an upright two-photon microscope (Sutter Instrument Company) equipped with an 8-kHz-resonant scanner (Cambridge Technology, Inc.), a pulsed femtosecond Ti:Sapphire laser (Chameleon Vision II, Coherent), a T565LPXR beam splitter (Chroma), ET525/70M and ET605/70M emission filters (Chroma), two GaAsP photomultiplier tubes (H10770PA-40 MOD; Hamamatsu) and a 16X 0.8 NA water-immersion objective (CF175; Nikon). Typical power used for imaging GCaMP6f in graft cells was 20-30 mW. Image resolution was 512 × 512 pixels at 30.95 frames per second.

Pinch stimuli were delivered with a rodent pincher system (2450, IITC Life Science, Inc.) and pressure sensor output was recorded using MCS software (Sutter Instrument Company; sampling rate, 1 kHz). Light touch and hindlimb movement through range of motion were applied manually by the experimenter over predetermined imaging frame acquisition numbers.

### Image Data Processing and Analysis

Slice imaging data were acquired as TIFF stacks and did not require motion correction because slices were held in place by a nylon harp. As imaging data acquisition began before Clampex protocols for excitation and stimulation were initiated, the frame at which the protocol began could be easily distinguished as the first frame with detectable fluorescence signal. A rapid period of photobleaching occurred as soon as the blue LED was turned on, so the first ten seconds after protocol initiation in each image stack were discarded to establish a smoother baseline fluorescence. When quantifying response amplitude or latency, image stacks were unfiltered. When quantifying activity dynamics, stacks were Kalman filtered using ImageJ software (Schneider et al., 2012). Regions of interest (ROIs) were manually drawn on the standard deviation projection of the stack around areas of fluorescence with neuronal morphology and dynamic fluorescence. These ROIs were converted to a mask and saved as a text image before being converted to a binary matrix in Matlab (Mathworks) with ones designating pixels belonging to ROIs. Image stacks with accompanying ROIs were then processed and analyzed with Matlab routines modified from a previously published calcium imaging toolbox (Romano et al., 2017). Briefly, background fluorescence from neuropil surrounding each ROI but not including neighboring ROIs was scaled by a factor of 0.9 and subtracted from the ROI’s fluorescence before the change in fluorescence over a slowly varying baseline (ΔF/F) was calculated for each ROI over the course of the imaging session. To quantify response strength, we took the mean ΔF/F over a 1 s period following stimulus onset. Response latency was defined as the temporal midpoint to peak fluorescence following stimulus onset, which corresponds to a time midway through the calcium spike.

Significant fluorescence transients were identified by their exceeding a threshold that was dynamically calculated based on the scale of the noise of the ROI’s fluorescence as well as the calcium reporter’s decay time constant. We then deviated from the Romano toolbox and took the first derivative of the ΔF/F trace, d(ΔF/F)/t, during these periods of significant fluorescence and subtracted 2.5 x (the standard deviation of d(ΔF/F)/t during non-significant periods). Those imaging frames with positive d(ΔF/F)/t values after noise subtraction were counted as periods during which true spiking activity took place within the cell, since the fluorescence decay period is not generally associated with activity (Carrillo-Reid et al., 2008). This data was used to generate raster plots of ROI activity.

Clusters of cells with similar activity dynamics were identified with scripts from the Romano toolbox based on the first derivative rasterization described above. Briefly, principal components analysis (PCA) was run on the z-scored activity of each ROI, and non-orthogonal factor rotation, or *promax* (Hendrickson and White, 1964), was applied to allow for non-exclusive cell assemblies, or clusters. A cell was designated as belonging to a cluster based on its z-scored loading on the principal component defining that cluster. A matching index was then used to quantify significant activations of the clusters over the imaging session, with the threshold p-value set at < 0.05 (Romano et al., 2017).

In vivo imaging data was converted from MDF format to TIFF stacks and corrected for lateral motion artifacts using TurboReg (Thévenaz et al., 1998). Uncut image stacks were Kalman filtered and ROIs were drawn around areas with dynamic fluorescence. For in vivo imaging only, ROIs included cell processes as well as soma, potentially allowing different regions of the same cell to be quantified separately. DeltaF/F traces were quantified as for slice image stacks, but with 0.4-factor or no background neuropil subtraction.

### Tissue Processing and Immunofluorescence

After imaging, slices were transferred to cold DaCSF and stored at 4°C until the end of the day and then transferred to 4% paraformaldehyde (PFA) in phosphate buffered saline (PBS). Slices were fixed for 20 minutes to overnight at 4°C and transferred to 30% sucrose in PBS for long-term storage. Following in vivo imaging experiments, animals were killed in their home cage using CO_2_ asphyxiation at a 20% fill rate, in accordance with IACUC guidelines. Animals were then transcardially perfused with PBS followed by PFA. The spinal column was dissected and placed in PFA at 4°C overnight, then transferred to 30% sucrose in PBS for long-term storage. Slices from ex vivo experiments were either sectioned at 15 µm on a cryostat (CM1950, Leica) and direct-mounted or stained unsectioned, free-floating. Spinal cords from in vivo experiments as well as whole brains were sectioned at 30 µm on a cryostat and stained free-floating. Standard staining techniques were used with primary antibodies against NeuN (guinea pig from Millipore, abn90, at 1:1000 to label neurons); green fluorescent protein (GFP, chicken from Aves, GFP-1020, or rabbit from Life Technologies, A6455, at 1:1000 to label GCaMP6f-expressing cells); and mCherry (goat from Sicgen, AB0040, at 1:1000 to label ChrimsonR-tdTomato). Secondary antibodies were all Alexa Fluor conjugates from donkey and included guinea pig 488 (Jackson Immunoresearch (JI), 706-545-148); goat 555 (Life Technologies (LT), A21432); chicken 488 (JI, 703-545-155); guinea pig 647 (JI, 706-605-148); and rabbit 488 (Invitrogen, A21206). For direct mount staining, primary and secondary antibody concentrations were quadrupled.

Stained sections were imaged on an automated all-in-one widefield microscope (BZ-X710, Keyence) or a manual widefield microscope (BX53, Olympus) with a Retiga 2000R Fast 1394 CCD camera (QImaging).

### Statistical Analysis

Two-group comparisons were tested by Welch’s t-test. Data are presented as mean ±SEM.

