## Supplemental Figures for "Calcium Imaging Reveals Host-Graft Synaptic Network Formation in Spinal Cord Injury"

#### Supplementary Figures

##### Figure S1: Identifying graft boundaries using differential interference contrast (DIC) optics

(A) DIC image of an intact spinal cord slice in the sagittal plane. (B) DIC image of a spinal cord slice in the sagittal plane after a dorsal column spinal cord lesion, with a neural progenitor graft (g) in the lesion site. The interface between host (h) and graft (g) is indicated by the sparsely dashed line. (C) Standard deviation projection of 50 sec calcium imaging video from the same field of view as in (B), showing the actual region of GCaMP6f fluorescence. The host/graft border is readily recognizable by DIC optics, particularly in the graft core. Expression of GCaMP6f is exclusively in grafted neurons because neural progenitor cells express Cre recombinase under the synapsin promoter, and these cells were mixed with a Cre-dependent AAV vector expressing GCaMP6f prior to grafting. Any AAV vector that might spread into the host could not express GCaMP6f because the host lacked Cre. In all images, the dorsal aspect of the slice is at the top and the rostral aspect is at left. Scale bars, 300 µm (A-C).

##### Figure S2: 4-AP increases graft spontaneous activity and response to corticospinal stimulation

(A) Light path for widefield, all-optical stimulation and imaging in acute slice prep. A 470 nm LED excites GCaMP6f for imaging neuronal activity, while a 617 nm LED stimulates regenerated corticospinal axons expressing Chrimson. LED1, 470 nm mounted LED; LED 2, 617 nm mounted LED; LPD, long-pass dichroic mirror; DM, deflecting mirror; BPD, band-pass dichroic mirror. Condensers and excitation and emission filters not shown for simplicity. (B,C) Standard deviation projection of GCaMP6f fluorescence in a neural progenitor cell graft, with regions of interest (ROIs) outlined in yellow in (C). ROIs were identified based on activity in calcium imaging videos and outlined on the standard deviation projection of the video to find the precise boundaries of dynamic GCaMP6f fluorescence. (D) Traces of the change in fluorescence over baseline of the 93 cells outlined in (C) in recording solution with no 4-AP added. Over the 50 sec shown, 4 trials of optogenetic corticospinal stimulation (vertical cyan bars) were performed at 10 sec intervals. Stimulation consisted of two 10 msec pulses from a 617 nm LED with a 50 msec start-to-start separation. Modest levels of spontaneous activity were observed. Red portions of traces indicate periods of significant activity. (E) As in (C), after wash-in of 100 µM 4-AP. More spontaneous activity was observed than without 4-AP, together with greater responses to corticospinal stimulation. (F) Quantification of mean activity (imaging frames with significant upward fluorescence transients (red portions of traces) divided by total frames (see Methods for neuronal activity deconvolution procedure)) before and after 4-AP wash-in in n = 123 cells in N = 3 animals during runs without stimulation. (G) Quantification of mean response strength to 8 trials of corticospinal stimulation (same protocol as in C-D, but with longer 500 msec train at 20Hz) before and after 4-AP wash-in in n = 53 cells in N = 3 animals. (H) Traces of change in fluorescence over baseline of 28 graft cells from an animal that did not receive AAV-Chrimson injections. While cells exhibited abundant spontaneous activity, no response to light stimulation was observed in N = 6 six animals with no AAV-Chrimson injections, indicating that the responses seen in graft neurons to optogenetic stimulation of regenerated corticospinal axons are not the result of direct photostimulation of non-opsin-expressing graft cells. In both images, the dorsal aspect of the slice is at the top and the rostral aspect is at left. Scale bar, 200 µm (B-C). Two-group comparisons were tested with Welch's t-test (**p<0.01). Data are represented as mean ±SEM.

**Figure S3: Graft neurons exhibit spontaneous activity in individual neurons and groups of simultaneously active neurons**

(A) Fluorescence traces of all graft neurons analyzed in (Figure 2) displaying varying frequencies and patterns of activity. Traces are ordered from lowest mean activity (top) to highest mean activity (bottom).

**Figure S4: Temporal sequence of calcium activity within assemblies**

(A) ΔF/F time series in 30 msec sequential epochs showing initiation of a spontaneous calcium transient in the magenta cluster, reaching peak activation over ~500 msec (see also Figure 2). The lower panel of images (B) shows the same cluster at higher brightness. The adjacent cyan cluster becomes active approximately 60 msec after the magenta cluster, raising the possibility that activity spreads from one cluster to another (or within a single, large cluster). However, frame-by-frame analysis of prolonged recordings of spontaneous activity did not confirm a repetition of spread from the magneta cluster to the cyan cluster. Most frequently, we observed simultaneous activation of most cells within single clusters, suggesting extensive interneuronal connectivity within neurons of single clusters, constituting localized neuronal assemblies. In all images, the dorsal aspect of the slice is at the top and the rostral aspect is at left. Scale bar, 200 µm.

**Figure S5: Graft neurons respond to corticospinal stimulation even in areas of low Chrimson expression**

(A) Example of a graft with three cells at the caudal end of the graft in the lesion site, outlined in yellow; these cells responded to corticospinal stimulation. Images are of native fluorescence, without antibody-mediated amplification of signal. (B) Higher magnification of inset from (A) shows a lack of detectable Chrimson-labeled corticospinal axons in the region of the responding cells. (C) Mean ±SEM shade traces of the responses of the three cells in (A-B) to 12 trials of the same optogenetic stimulation (vertical cyan bar). (D) Overlay of response strength in graft cells on native ChrimsonR-tdTomato fluorescence showing responsive cells in areas of both high and low Chrimson brightness. (E) No correlation was seen between the response strength and Chrimson brightness in n=68 cells from N=3 animals. Response strength and Chrimson brightness were normalized to the respective maximums in each field of view. A least-squares linear fit to the data is shown. In all images, the dorsal aspect of the slice is at the top and the rostral aspect is at left. Scale bars, 300 µm (A); 150 µm (B, D).

**Figure S6: Spontaneous, correlated calcium activation occurs across entire field-of-view in intact dorsal horn**

(A) Frame-by-frame ΔF/F time series showing initiation of a calcium transient uniformly across the entire field-of-view in intact sagittal spinal cord dorsal horn. In all images, the dorsal aspect of the slice is at the top and the rostral aspect is at left. Scale bar, 200 µm.

**Figure S7: Optogenetic stimulation of corticospinal axons activates individual neurons and neuron clusters in intact host spinal cord**

(A) Fluorescence traces of host neurons showing spontaneous activity and responses to 500 msec optogenetic stimulation of corticospinal axons. (B) Mean response traces with ±SEM shade of n = 10 cells in one slice to four trials of corticospinal stimulation. (C) Standard deviation projection of the run shown in (A) with all ROIs outlined in yellow and assemblies of neurons with similar activity dynamics indicated by fill color. (D) First derivative raster plot (top) of traces from (A), with cells grouped into the assemblies shown in (C). Cells above red line belonged to assemblies. Significant activations of color-coded assemblies are displayed below. In (C), the dorsal aspect of the slice is at the top and the rostral aspect is at left. Scale bar, 200 µm.

#### Figure S1

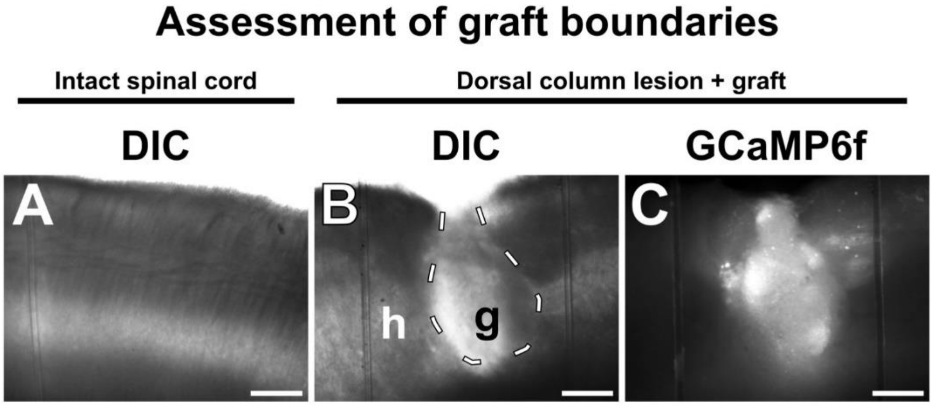
Assessment of graft boundaries
Intact spinal cord
Dorsal column lesion + graft
DIC
DIC
GCaMP6f
A
B
C
h
g

### Figure S2

#### Acute slice calcium imaging

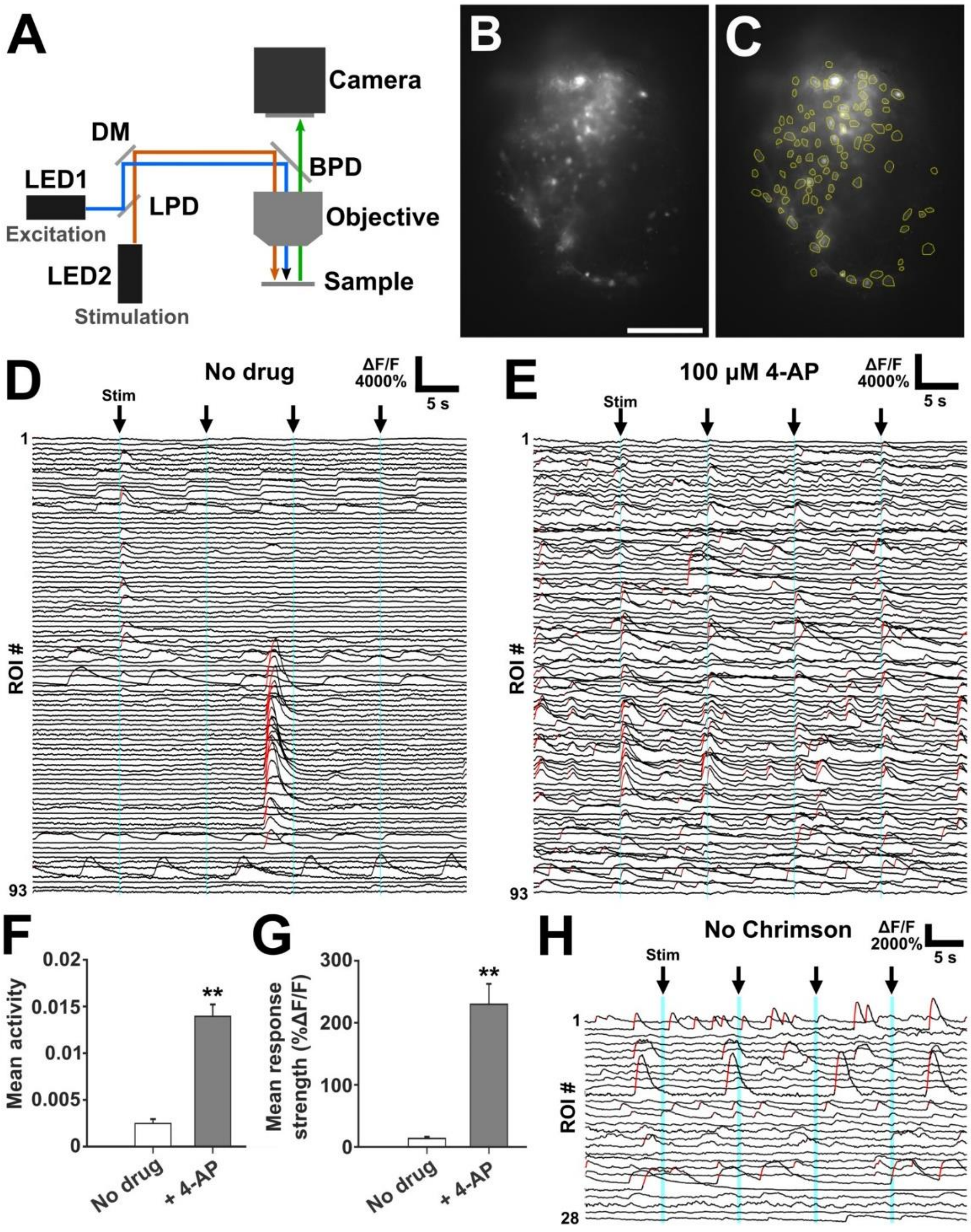

Figure S3

#### Spontaneous graft activity

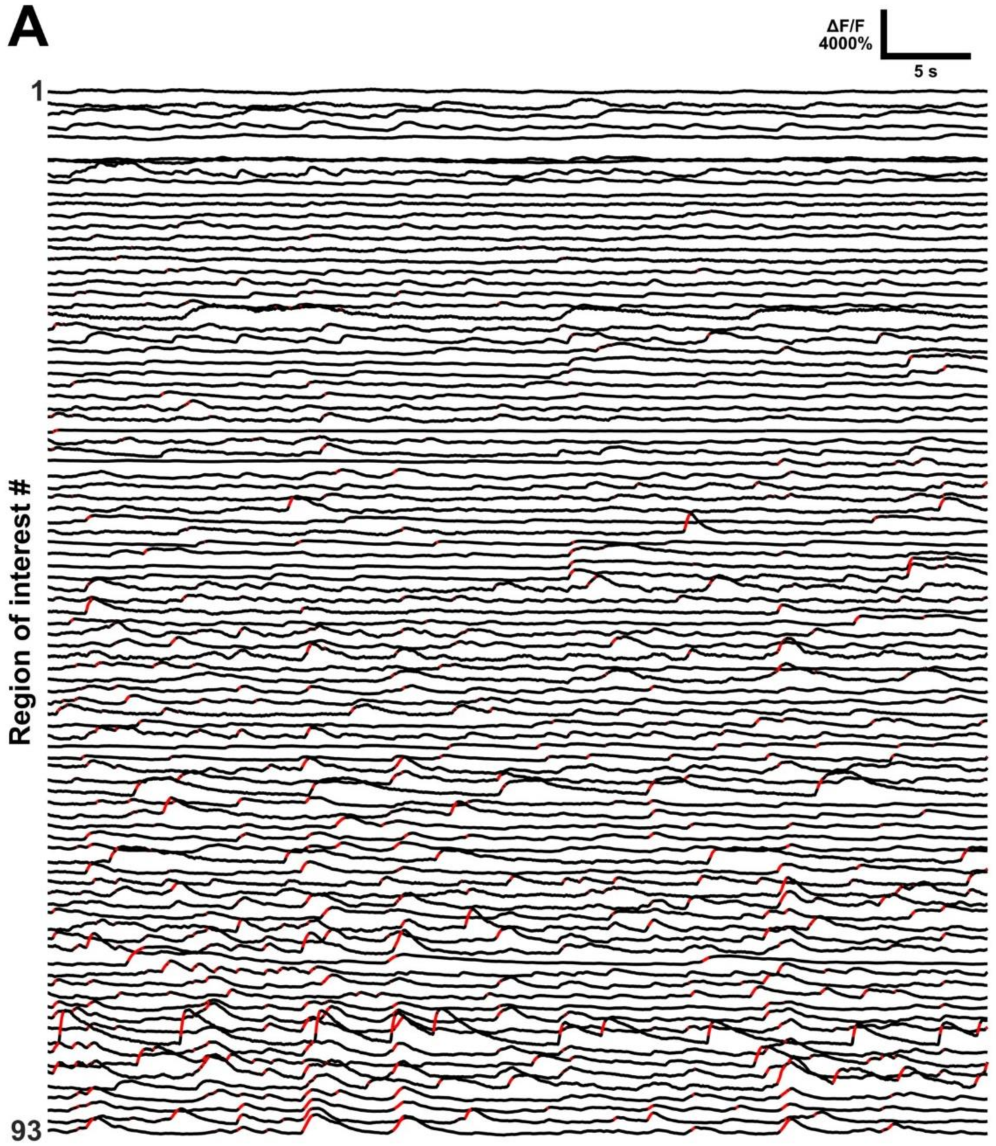

#### Figure S4

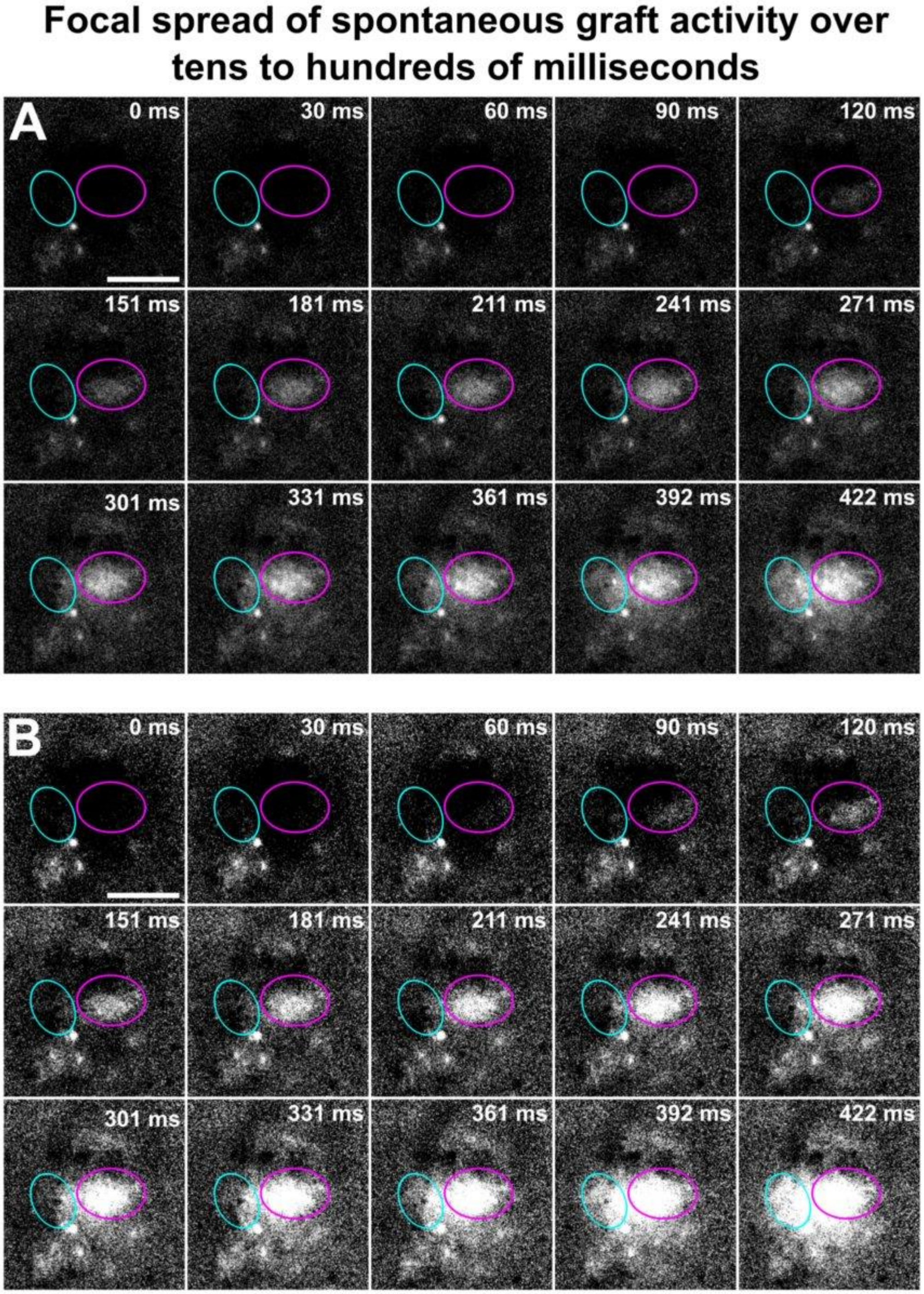
Focal spread of spontaneous graft activity over tens to hundreds of milliseconds
A
0 ms
30 ms
60 ms
90 ms
120 ms
151 ms
181 ms
211 ms
241 ms
271 ms
301 ms
331 ms
361 ms
392 ms
422 ms
B
0 ms
30 ms
60 ms
90 ms
120 ms
151 ms
181 ms
211 ms
241 ms
271 ms
301 ms
331 ms
361 ms
392 ms
422 ms

### Figure S5

#### Caudal graft responds to CST stimulation

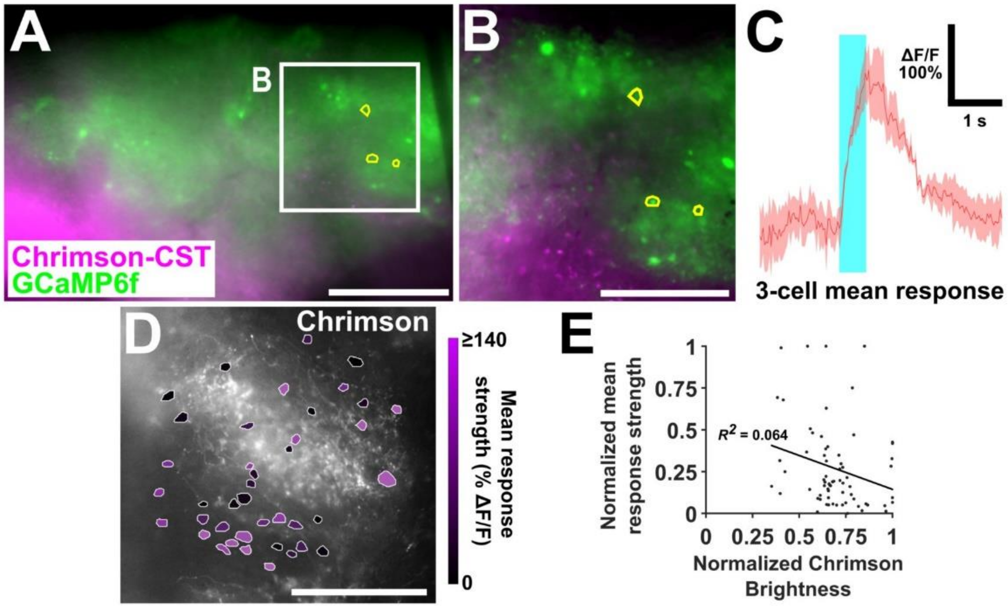

#### Figure S6

### Intact spinal cord neuron assemblies are activated uniformly

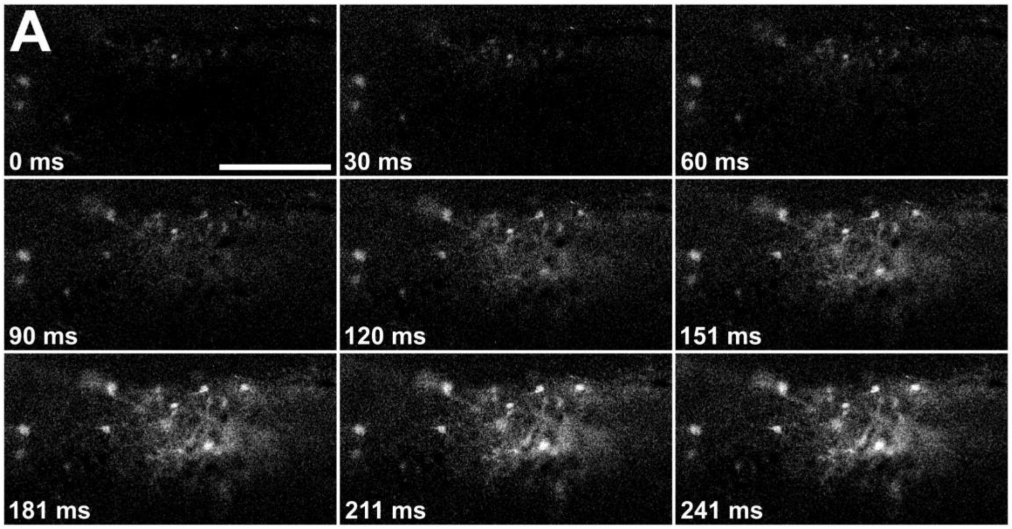

#### Figure S7

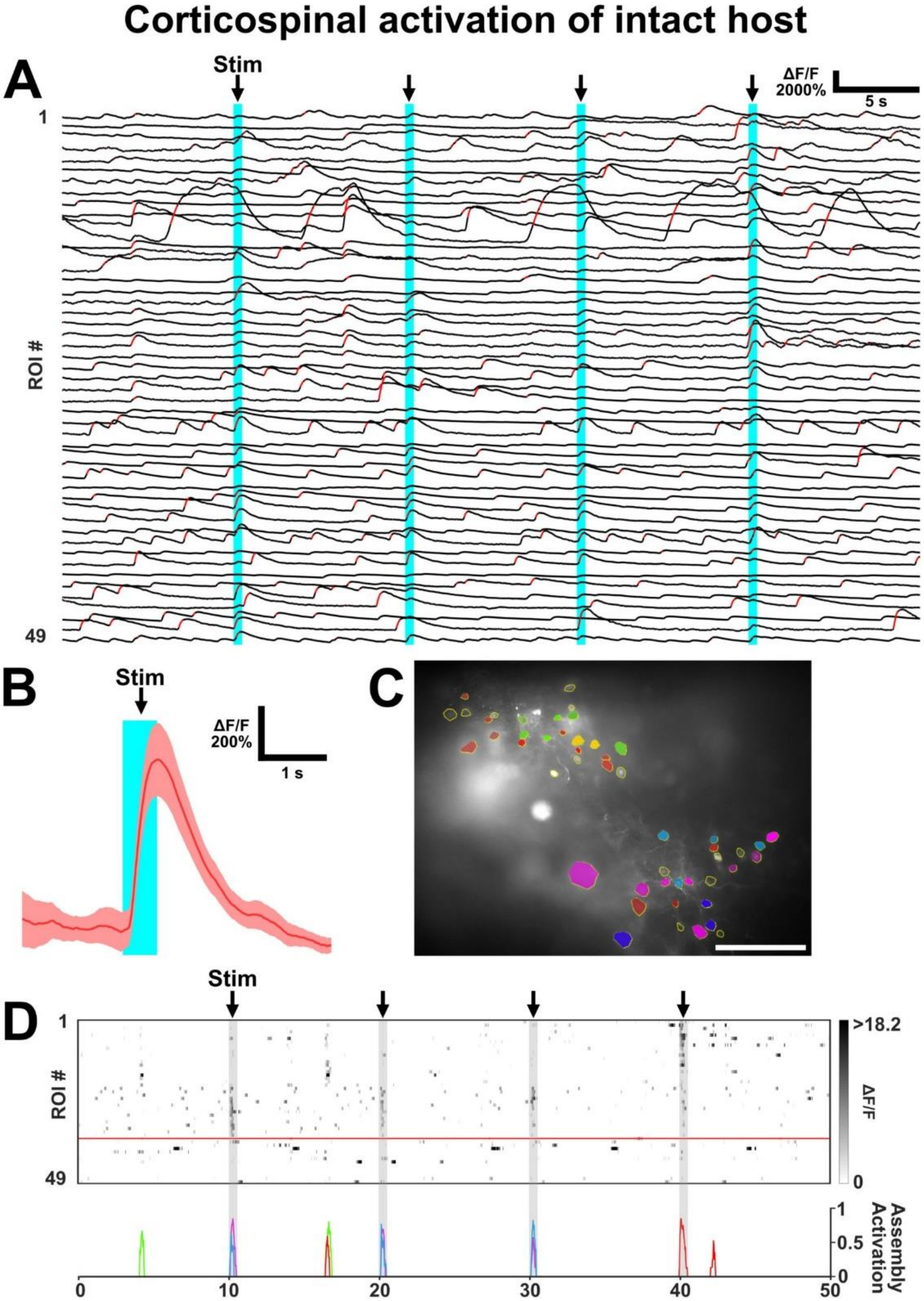

Corticospinal activation of intact host
A
Stim
ΔF/F
2000%
5 s
1
ROI #
49
B
Stim
ΔF/F
200%
1 s
C
D
Stim
1
ROI #
49
>18.2
ΔF/F
0
1
0.5
0
Assembly Activation
0
10
20
30
40
50
